# Longitudinal Tracking Reveals Developmental Transitions in Zebrafish Clock Gene Expression

**DOI:** 10.64898/2026.05.18.726011

**Authors:** Camila Morales Fénero, Raina E. Sacksteder, Jacqueline M. Kimmey

**Affiliations:** Department of Microbiology and Environmental Toxicology, University of California, Santa Cruz; Santa Cruz, USA

## Abstract

Circadian clocks coordinate physiological and behavioral rhythms by synchronizing biological processes with environmental cues. These rhythms emerge during development, but it remains unclear whether their component genes are activated by a common program or assembled through distinct regulatory pathways. To address this, we used longitudinal luciferase reporters to monitor *per3* and *per2* expression across zebrafish embryonic and larval development. Although both genes are canonical components of the circadian clock, they showed strikingly different developmental regulation. Two temporal frames of circadian gene expression were identified: an embryonic stage and a larval stage, each evident under different entrainment conditions. *Per3* displayed early rhythmic expression in light/dark conditions, which was independent of *per2 and cry1a* light-entrainment regulation, but required *bmal* activity. Meanwhile, *per2* displayed light-responsive transcription and remained largely *bmal*-independent. At the same time, both genes exhibited an endogenous embryonic expression that could not be explained solely by light-driven regulation, indicating that developmental inputs contribute to clock gene activation before mature larval rhythms are established. These findings demonstrate that the zebrafish circadian system is not assembled through a single synchronized onset of clock gene expression, but through gene-specific regulatory programs that shift across development.

## Introduction

Biological processes across living organisms are controlled by an intrinsic timekeeping mechanism known as the circadian clock. This molecular oscillator generates 24-hour rhythms that coordinate physiology, metabolism, and behavior in anticipation of daily environmental cycles^1^. The circadian clock in vertebrates operates through a transcriptional–translational feedback loop (TTFL): the transcription factors CLOCK and BMAL1 heterodimerize and activate the expression of Period (Per) and Cryptochrome (Cry) genes via E-box enhancer elements in their promoters. PER and CRY proteins subsequently accumulate, translocate to the nucleus, and repress CLOCK:BMAL1 activity, thereby inhibiting their own transcription^2^. Auxiliary feedback loops involving nuclear receptors such as REV-ERBα/β and RORα further stabilize the core loop, providing robustness and precision to the oscillation^3^.

Despite decades of research into the molecular mechanisms of the circadian clock, a central question remains: when during development does a functional oscillator first emerge, and what regulatory forces shape its onset? Looking across different organisms, a similar picture appears: an endogenous expression of clock gene precedes the onset of rhythmicity. In mammals, circadian clock oscillations emerge gradually during mid-to-late gestation while clock gene components in the mouse master regulator, the suprachiasmatic nucleus (SCN), are expressed as early as embryonic day 12–15^4,5^. In *Drosophila, Per* expression in the presumptive central-clock dorsal neurons begin to oscillate in the embryo in constant darkness, and a single 12-hour light pulse during the embryonic stage is sufficient to synchronize adult activity rhythms^6^. This implies that both the oscillator and its entrainment machinery are established early in development, however, the regulatory mechanisms that drive the developmental onset of clock gene expression in vertebrates remain poorly understood. The zebrafish embryo has emerged as a powerful model for studying the developmental emergence of the circadian clock. Large clutches of optically transparent embryos develop *ex-utero* in few days, with high genetic tractability and a circadian clock conserved with other vertebrates. Studies in the early 2000, showed that zebrafish clock genes have an early onset of expression in development^7–9^. Although the central activators *clock1* and *bmal1* do not oscillate until 4 days post-fertilization (dpf) in light: dark (LD) conditions, rhythmic expression of *per1, per2* and *per3* have staggered onset during development, with detection as early as ∼20 hours post-fertilization (hpf) ^7–9^. These studies have given initial insights into zebrafish circadian clock development, but due to the technological constraints of the period, this was achieved with endpoint analysis of gene expression across multiple animals and timepoints. Nowadays, the generation of luciferase transcriptional reporters enables continuous monitoring of gene expression in the same animal over extended periods. However, there are no current reports analyzing longitudinal expression of circadian clock genes during zebrafish development. Here, we address this gap by using *per3:luc* and *per2:luc* reporter lines to characterize the dynamic expression of these two clock genes continuously from 0 to 7 dpf and explore the regulatory mechanisms behind their expression.

## Results

### Circadian clock genes show differential windows of expression during development

We began by analyzing *per3:luc* in embryos from hemizygous adults whose spawning of egg was settled as ZT0 and synchronized with lights on so that ZT0 corresponds with 0 hours post-fertilization (hpf). *Per3:luc* developing embryos exposed to 12h:12h light: dark (LD) cycles for 7 days displayed oscillatory expression with a peak at ZT6 and a period of ∼24h (Fig.1A, blue line). The cycling expression of *per3* was detected as early as 12 hpf (eJKT p>0.05) (Fig.1A, insert). This was visualized again with an inverted LD cycle (DL) of 12h dark and 12h light, where *per3* expression showed robust expression in phase with light (Fig.1A, black line). Although *per3* has not been described as a light inducible gene, its expression pattern shifted rapidly under inverted light conditions, prompting us to compare its patterns with *per2*, a known light-induced gene^10–13^. Unlike *per3*, expression of luciferase under the *per2* promoter continuously increased in a staggered fashion, without returning to baseline levels during dark periods, peaking at the third day of development (Fig.1B). Light exposure continually increased *per2* expression after 24 hpf, independently of the LD condition used (Fig.1B). Moreover, *per2* expression kinetics changed after 96 hpf with a rapid increase with a peak at ZT4 and a slow decay until the next light exposure. This supports the idea that *per3* and *per2* are regulated by different pathways.

**Figure 1.**
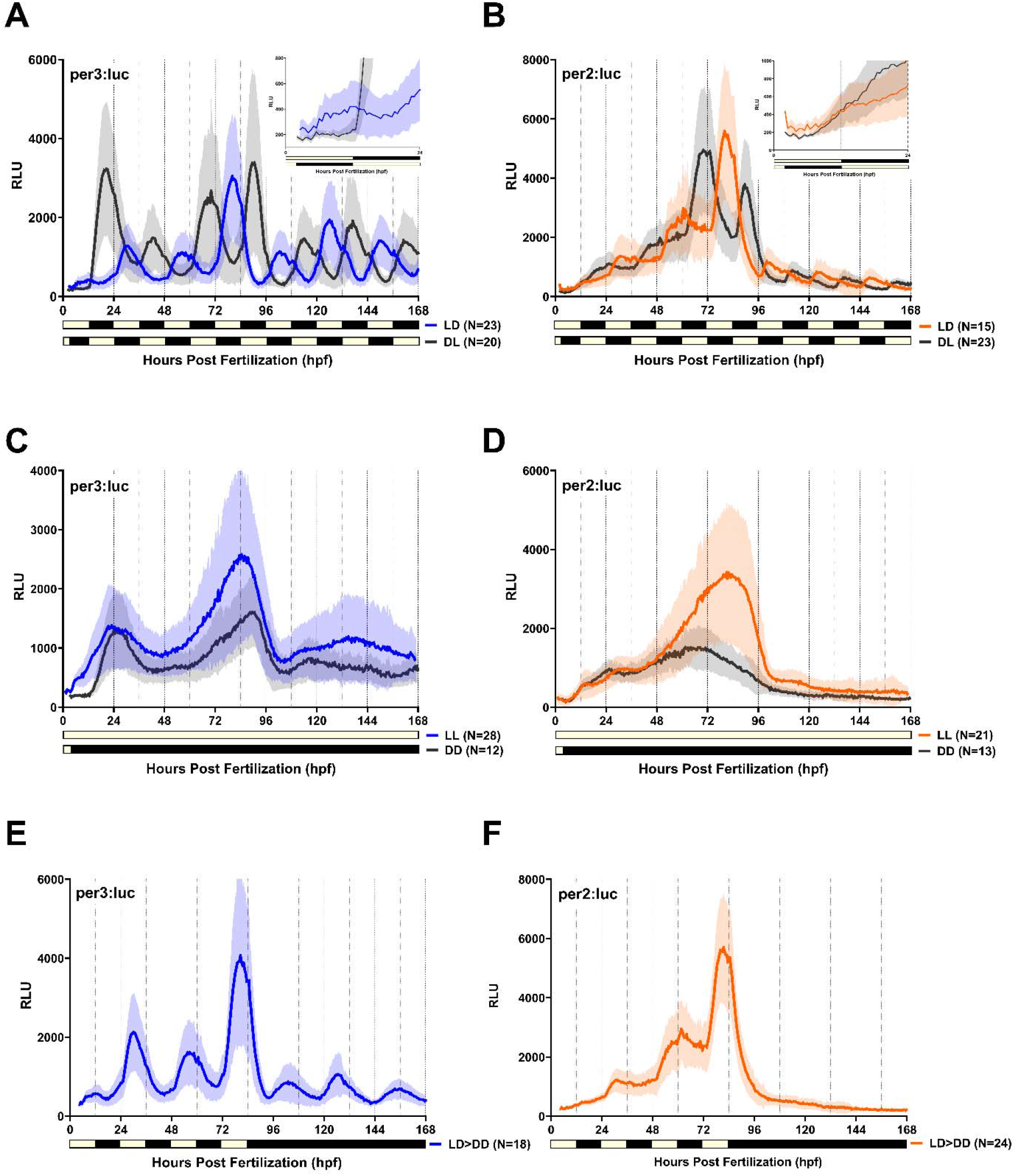
Zebrafish clock genes display two windows of expression during development. Luminescence reads from *per3:luc* (left) and *per2:luc* (right) zebrafish embryos of from 0 to 168 hours (7 days) of development in different light settings: **A-B**. 12h:12h cycles of Light:Dark (LD) or Dark:Light (DL), **C-D**. Constant Light (LL) or Constant Darkness (DD), and **E-F**. 12h:12h Light:Dark entrainment for 4 days into Constant Darkness (LD>DD). RLU = Relative Luminescence Units. N=Number of animals per group.

Nonetheless, we noticed a common feature on both genes, where there is high amplitude expression in the first 4 days of development that then decreased and stabilized after this period. We therefore tracked the developmental expression of *per3* and *per2* in the absence of LD entrainment to verify basal expression without the interfering of light. To our surprise, in constant darkness *per3* presented a double peak expression with an infradian rhythm of a ∼29-hour period (BD2-Classic p>0.05) (Fig.1C; black line). After 96 hpf, expression was variable, with a low amplitude pattern that showed ∼24h rhythmicity in 75% of animals (eJTK p>0.05). This indicates that there is a basal or developmental expression of *per3* that does not rely on external stimuli. In constant light, *per3* showed similar expression patterns as in constant darkness, suggesting that its rhythmic expression is more responsive to oscillating lighting conditions rather than light itself (Fig.1C, blue line). In the case of *per2*, animals in constant darkness showed no obvious oscillatory patterns of expression other than a relative peak at ∼60 hpf, after which it declined. In constant light conditions a sharp divergence is seen starting from 48 hpf, with a marked increase in amplitude and a peak shifting to approximately 84 hpf (Fig.1D). After 96 hpf, *per2* expression flattened in both constant light and dark conditions. Although *per2* has been generally described as a light-driven gene^11^, it has shown tissue-specific developmental expression independent of light^8^, suggesting that its transcription may be regulated by different pathways depending on the stage of development.

One key characteristic of circadian regulated expression is the ability to sustain autonomous cycles in constant conditions or free run^14^. We therefore analyzed *per3* and *per2* expression in these settings. In line with previous reports^10^, *per3* oscillations persisted in constant darkness after 4 days of entrainment (Fig.1E). This was also seen in 4XE-box:luc animals which express luciferase under an artificial promoter with 4 tandem E-box sequences, the binding motif for Clock:Bmal^15,16^ (Fig.S1). Interestingly, the E-box reporter expression has a peak ∼6h earlier than *per3*, in the expected transition from dark to light (ZT24/0). This suggests that the E-box reporter may reflect direct Clock:Bmal activity while *per3* transcription may be delayed by other regulatory factors. In contrast, *per2* did not show expression in the absence of light after 4 dpf (Fig.1F), indicating that *per2* regulation is exclusively light inducible during the larval period. Together, these data depict specific developmental windows of expression for both clock genes: An embryonic window from 0 to 4 dpf, and a larval window from 4 dpf to (*at least)* 7 dpf. The basal expression of *per3* and *per2* revealed in constant conditions also suggests a light independent endogenous regulation during the embryonic stage, that can be concomitant with the circadian/light-controlled regulation of the clock. Moreover, the aligned peaks of expression of *per3* and *per2* during LD cycles suggest that a light-dependent mechanism synchronizes their expression.

### Light entrainment of per3 is independent of per2 and cry1a regulation

Consistent reports have shown that light can directly activate the rapid transcription of genes in the zebrafish, including the two clock genes *per2* and *cry1a*^12,13^. The current model shows that these genes are activated through D-boxes located in their promoters^11,17,18^. Although *per3* can sustain oscillations in constant darkness, like the 4XE-box:luc reporter (Fig.1 and S1), its expression is directly affected by changes in lighting conditions. *Per3* does not contain a canonical D-box sequence in its promoter ^10,19^, therefore, the light-dependent effects we observe are likely to arise indirectly, potentially through light-driven effects on the core oscillator machinery. *Per2* and *cry1a* have been previously implicated in the regulation of behavioral entrainment in zebrafish larvae but their role in light regulation of the molecular feedback loop is less clear^13,20–22^. Using CRISPR-Cas9 gene editing, we generated double knockout (DKO) crispants of the light-inducible genes (LIG) *per2* and *cry1a* (hereafter LIG DKO) and analyzed their effect on *per3* expression. We verified that tyrosinase-knockout (tyr KO) animals did not differ from uninjected controls and therefore used uninjected animals for the remainder of the experiments (Fig. S2). Under LD exposure, *per3* expression showed no difference between uninjected controls and LIG DKO animals from 0-4 dpf, although a small shift was apparent after 4 dpf in LIG DKO animals (Fig.2A). However, inverted light cycles did not show any differences between control and DKO groups (Fig.2B). We confirmed the presence of target mutations in injected animals using the T7 Endonuclease I system which detects mismatches caused by CRISPR-induced indels. To ensure that *per3* can still respond to light changes in the absence of *per2/cry1a* we submitted the animals to 4 days of LD entrainment, followed by a 12-hour shift in the light cycle and continuous darkness. Oscillations of uninjected controls and LIG DKO *per3:luc* animals rapidly responded to the light-induced shift and continued to cycle with a peak belonging to the last light period (Fig.2C). These results suggest that *per2* and *cry1a* are not required for *per3* light entrainment and that other unknown light regulatory mechanisms are involved.

**Figure 2.**
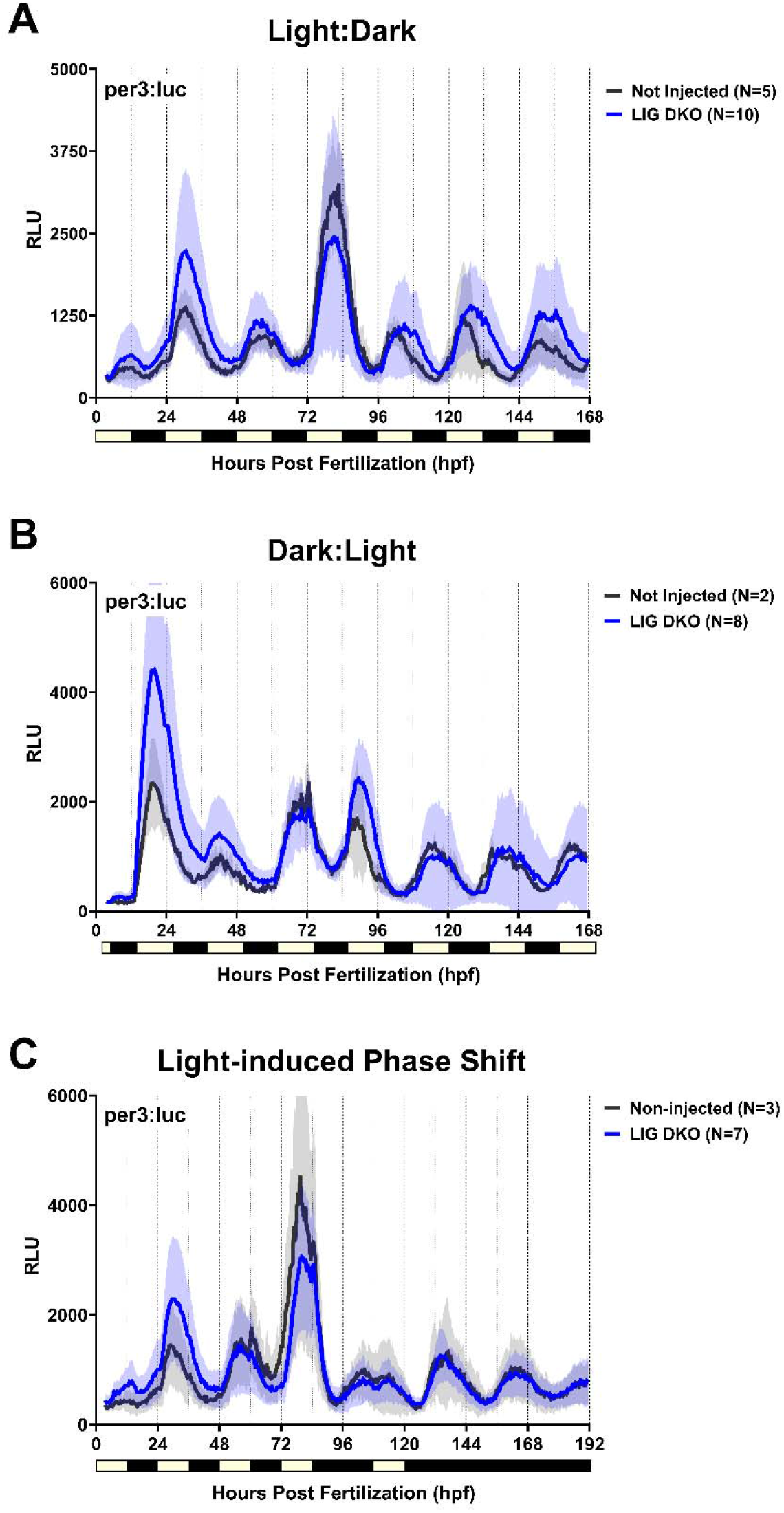
The light-induced genes *per2* and *cry1a* are not required for *per3* light entrainment. Luminescence reads from *per3:luc* embryos injected at 1-cell stage with a ribonucleoprotein (RNP) complex of Cas9 protein and gRNAs targeting the light-induced genes (LIG) *per2* and *cry1a* to generate double knockout (DKO) crispants. Controls are uninjected embryos. **A**. Tracking of luminescence of embryos across 7 days of development in **(A)** 12h:12h LD cycles, **(B)** 12h:12h of Dark:Light cycles and **(C)** 12h:12h of Light:Dark cycles for 4 days and 12h of an induced light phase shift followed by constant darkness. RLU = Relative Luminescence Units. N= Number of animals per group.

### Bmal is required for per3 transcription but not for per2 expression

A central molecular driver of circadian rhythmicity in vertebrates is the transcriptional-translational feedback loop in which CLOCK and BMAL drive transcriptional activation of *per* and *cry*, and PER and CRY proteins in turn repress CLOCK:BMAL activity. The components of the molecular clock are conserved in zebrafish, although the specific contributions of the duplicated clock gene paralogs remain incompletely understood. Using a dominant-negative version of *Clock1* previous studies have shown that Clock is required for the rhythmic onset of *per1* developmental expression^9^. We decided to test the role of *bmal* in the developmental expression of *per3* and *per2*, using the same crispants approach from the previous section. Initial experiments targeting the single and double paralogs *bmal1a, bmal1b* and *bmal2* individually showed no or low effect on *per3* expression, suggesting redundancy in their function (Fig.S3). We next generated a *bmal* triple knockout (*bmal* TKO) and analyzed the developmental expression of *per3:luc* in these animals. In LD settings *per3:luc; bmal* TKO animals lost their rhythmic expression over time, with a more dramatic effect after 4 dpf (Fig.3A). Similarly, *per3* expression in constant darkness showed a decrease in luminesce reads to completely flat expression after 4 dpf (Fig.3B). These results indicate that *per3* regulation requires *bmal* for its transcriptional activation and oscillatory nature.

**Figure 3.**
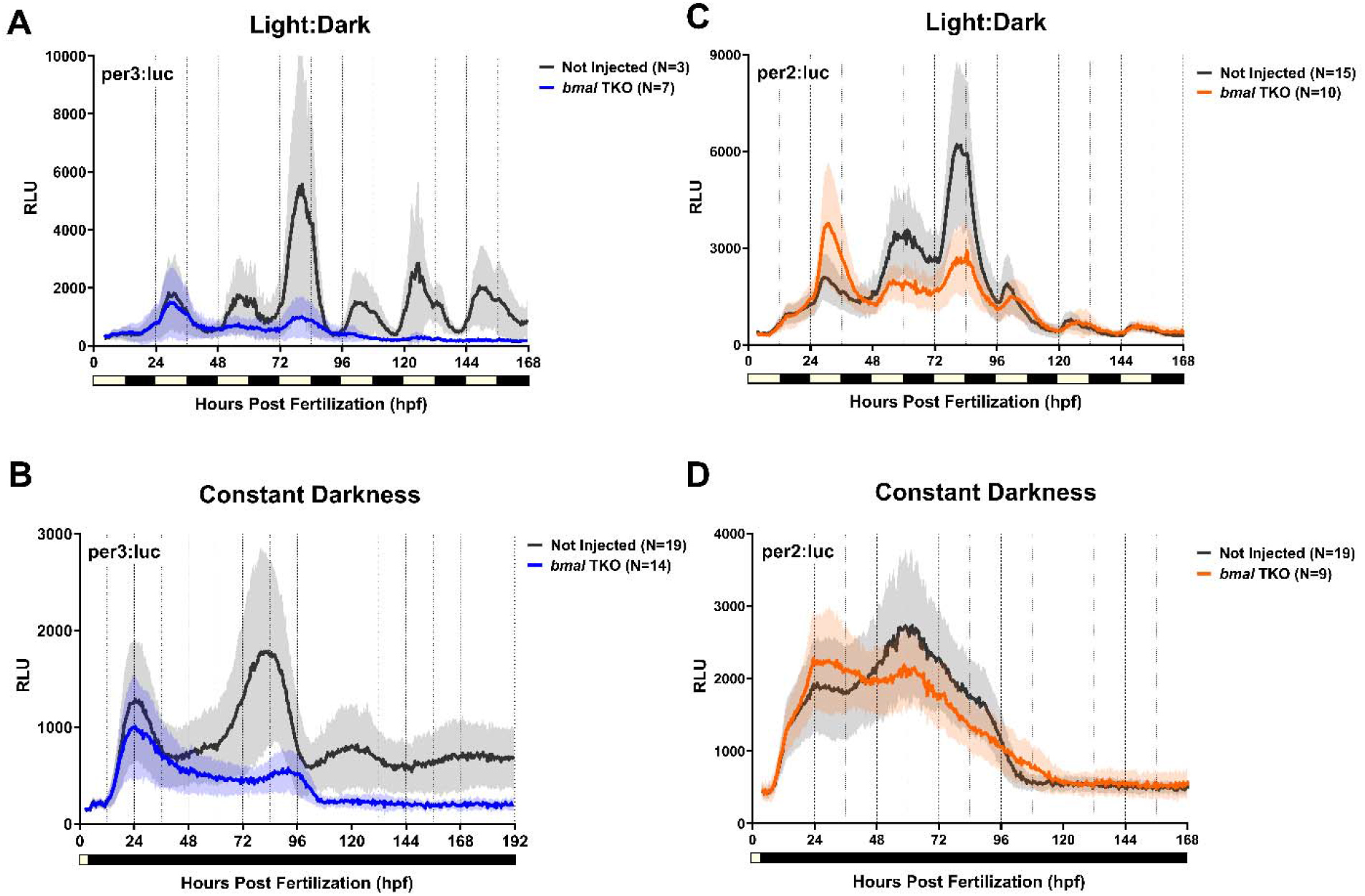
Bmal is essential for *per3* but not *per2* expression. Luminescence reads from *per3:luc* (left) and *per2:luc* (right) embryos injected at 1-cell stage with a RNP complex of Cas9 protein and gRNAs targeting the three zebrafish *bmal* genes: *bmal1a, bmal1b* and *bmal2*, to generate triple knockouts (TKO) crispants. Controls are uninjected embryos. Tracking of luminescence of embryos across 7 days of development in **(A, C)** 12h:12h LD cycles and **(B, D)** constant darkness. RLU = Relative Luminescence Units. N= Number of animals per group.

Interestingly, under LD conditions, *per2* remained light responsive in the absence of *bmal*, although its developmental expression profile was modified, with higher expression early and reduced peak amplitude later in development (Fig. 3C). Likewise, *per2* embryonic expression seen in constant darkness was slightly reduced but not totally lost (Fig.3D), suggesting that *bmal* may have a secondary regulatory role in *per2* expression rather than be the main activator of transcription. Unfortunately, the transcriptional programs responsible for *per2* embryonic activation are still unknown and will be focus of future studies.

## Discussion

A central question in chronobiology is understanding when during development a functional circadian clock first emerges and what mechanisms are behind this process. In this study, we took advantage of the zebrafish *ex-utero* development to try to answer this question. Using available transcriptional reporters expressing luciferase under the protomer of *per3* and *per2* genes, we characterized longitudinal expression patterns from early embryogenesis through larval stage. Our data revealed three main findings with important implications for the understanding of circadian clock regulation and development. First, both *per3* and *per2* exhibit two temporally distinct windows of expression, an embryonic phase (0-4 dpf) and a larval phase (4-7 dpf) with an endogenous light-independent component during the embryonic period. Second, our data shows that the light-inducible genes *per2* and *cry1a* are dispensable for light-mediated entrainment of *per3*, implying the existence of an uncharacterized light input mechanism. Third, the *bmal* paralogs are essential for *per3* developmental and oscillatory expression but play only a modulatory effect on *per2* regulation. Together, our results provide a new temporal map of zebrafish circadian clock expression during development and amplify the understanding of regulatory pathways that shape clock gene expression during ontogeny.

Our longitudinal analysis of *per3* and *per2* reporters extends and refine previous endpoint studies that described differential onsets of these genes, where rhythmic *per3* expression precedes that of *per2* during development^8^. Our approach reveals fundamentally different expression kinetics between *per3* and *per2*, with the presence of an endogenous light-independent expression during the embryonic window. This endogenous double-peak expression of *per3* in constant conditions and the light-independent embryonic expression of *per2* could be explained by tissue specific expression during development^7,8^. Previous reports have shown rhythmic expression of *per3* strongly concentrated in the retina and the pineal gland from 40 hpf^7,8^. Meanwhile, endogenous *per2* expression has been detected in the olfactory bulb and pituitary from 36 hpf^8^. However, no studies have been conducted analyzing the tissue expression of these genes at earlier timepoints. Consequently, the assessment of *per2* and *per3* transcripts expression in the whole embryo at the peak timepoints revealed in this study could give us a better spatial understanding of this process. Moreover, the mechanisms of activation of endogenous *per3* expression could spring from residual maternally inherited circadian transcription factors in the embryo. Numerous studies have identified accumulation of maternal mRNA of clock genes in oocytes and 1-cell stage embryos that persist until gastrulation stages^7,9,10^, however, no studies have explored the transfer and turnover of circadian maternal proteins in zebrafish embryos. In the case of *per2*, studies have shown Roraa light-independent activation through binding to ROR response elements (RORE) in the promoter of this gene^23^. Interestingly, *roraa* is expressed in retina and brain during development^24,25^. The role of *roraa* in the regulation of developmental *per2* will be the subject of future investigations.

The stabilization of clock gene expression after 4 dpf in the zebrafish is reminiscent of the gradual developmental maturation of behavioral circadian rhythmicity over the first days of life reported by Hurd and Cahill. Here, embryos deprived from entrainment cues early in development showed less rhythmic activity than embryos entrained for 4 or more days^26^. Dekens & Whitmore also mentioned this stabilization with the discovery of non-rhythmic expression of the core circadian activators *clock1* and *bmal1* in light entrainment conditions until the 4 day of development, where they become rhythmic^9^. However, repressor genes like *per1* showed rhythmic expression from 21 hpf, which was lost in embryos injected with a dominant-negative version of *Clock1*. These results, together with our data showing loss of *per3* oscillations in *bmal* TKO fish, suggest that core clock proteins are active during the embryonic phase and are required for the rhythmic expression of *period* genes in entrainment conditions, possibly through post-translational regulation. We speculate that the embryonic window may represent a period of clock maturation and calibration, during which light is required to consolidate and amplify emerging oscillations rather than initiate them. This idea is supported by *nr1d1* single-cell reporter data demonstrating that light is required for the amplitude and robustness of circadian oscillations in the developing zebrafish brain^27^. Notably, constant light enhanced the amplitude of both *per3* and *per2* expression in our experiments, suggesting that while light is not strictly required for embryonic clock gene expression, it serves as a potent amplifier of the emerging oscillatory machinery.

The canonical model of light-driven gene expression in zebrafish centers on the D-box enhancer as the molecular pathway through which light signals activate transcription of target genes including *per2* and *cry1a*^11,17,18^. Because loss of *per2* and *cry1a* have shown to impair behavioral rhythms in zebrafish larvae, they have been proposed as essential mediators of clock light entrainment^13,20–22^. Our results instead suggest that this behavioral phenotype may reflect impaired downstream outputs, as genetic ablation of both *per2* and *cry1a* did not disrupt light entrainment of *per3*, indicating that the molecular oscillator remains light responsive in their absence. The ability of *per3* to rapidly re-entrain to a shifted light schedule in the absence of *per2* and *cry1a* is particularly interesting, as it demonstrates that the light-responsive pathway controlling *per3* phase is functionally intact and independent of these light-induced genes. This result implies the existence of an uncharacterized mechanism by which light information reaches and resets E-box–dependent transcription. This pathway may operate through post-translational modifications, indirect transcriptional regulation of Clock:Bmal activity, or through alternative photosensory cascades that remain to be identified

The transcriptional activator complex Clock:Bmal is considered the master driver of *per* and *cry* gene expression in the vertebrate circadian clock. Here, we confirmed that this stands for *bmal* function regarding the oscillatory expression of *per3* but revealed a likely secondary role for the regulation of *per2*. The fact that *per3* expression declines over time in the *bmal* TKO may be a limitation of the crispants methodology. Crispants are primary injected and therefore mosaics. Phenotypic effects are not evident immediately as mutations accumulate over time, being apparent around 1-3 dpf, depending on the targeted gene^28–30^. Moreover, the embryonic period may be partially buffered from loss of Bmal function by maternal clock proteins, therefore we are not able to see the complete loss effects until maternal proteins are completely degraded^31,32^. On the other hand, light-induced *per2* expression was preserved, but with only slight impacts on its endogenous embryonic expression. These results indicate that Bmal is not one of the primary drivers of *per2* transcription. The nature of the endogenous activators of *per2* remains unknown, but candidates might include developmental transcription factors or other clock activators such as Roraa, that are active during development^23–25^.

This study provides the first comprehensive longitudinal characterization of circadian clock gene expression during zebrafish embryonic and larval development. We defined two temporal windows of expression in two clock genes with distinct regulatory mechanisms. We discarded the role of a known light input pathway in *per3* entrainment and revealed a fundamental difference in the Bmal-dependence of *per3* versus *per2* transcription. These findings reframe our understanding of how the vertebrate circadian clock is assembled and entrained during development.

## Methods

### Animals

Zebrafish husbandry and experiments were conducted in compliance with guidelines from the U.S. National Institutes of Health. All experiments were conducted under a protocol approved by University of California Santa Cruz Institutional Animal Care and Use Committee (IACUC) and with an approved Biohazardous Use Authorization. Adult zebrafish were raised in a 14 hour light, 10 hour dark lighting cycle at 28℃ and fed 2-3 times per day. Male and female adults were paired in mating tanks in the afternoon and fertilized eggs were collected the next morning after lights on. Eggs were transferred into petri dishes containing E3 medium (5 mM NaCl, 0.17 mM KCl, 0.33 mM CaCl2) and raised at 28°C. Animals used here include wild type AB, transgenic zebrafish carrying luciferase under a 26kb promoter of *per3 (per3:luc)*^10^, 1.9 kb promoter of *per2 (per2:luc)*, or the artificial tandem E-box promoter^33^ (4XE-box:luc).

### Cloning of the per2:luciferase construct

Gateway Cloning using the Tol2kit v1.0 was used to clone the zebrafish per2 promoter upstream of the Luciferase gene. The cloned region of the per2 promoter was based on Vatine et al., 2009 [^11^]. A fragment containing 1,813 bp of the 5′ flanking region and 164 bp of the 5′ untranslated region (UTR) of the per2 gene (accession number FJ435339.1) was PCR amplified from genomic DNA using specific primers containing attB sites: per2 forward, 5’-GGGGACAACTTTGTATAGAAAAGTTGCTATCATTTCCCAGTGCTTAGTGGC-3’, and per2 reverse, 5’-GGGGACTGCTTTTTTGTACAAACTTGACTGACAACTTCAGCAAATCTTCTT-3’. The PCR product was validated using gel electrophoresis, and a BP reaction was carried out using Gateway BP Clonase II (Invitrogen) to insert the per2 promoter into the pDonorP4-P1r vector according to the manufacturer’s protocol and create the per2 5’ entry vector. The construct was transformed into competent Top10 E. coli cells, purified, and validated using a SacI-HF (NEB) digest. A Luciferase middle entry vector was cloned from the 4xE-box:Luc plasmid kindly provided by Thomas Dickmeis, and is described in Weger et al., 2013 [^33^]. Primers containing attB sites were used to amplify the region containing the Luciferase gene from the 4xE-box:Luc plasmid: luciferase forward, 5’-GGGGACAAGTTTGTACAAAAAAGCAGGCTTAACAGCGGAGACTCTAGAGGG-3’, and luciferase reverse, 5’-GGGGACCACTTTGTACAAGAAAGCTGGGTCATCGCTGAATACAGTTACATT-3’. Again, the PCR product was validated using gel electrophoresis, and a BP reaction was carried out using Gateway BP Clonase II (Invitrogen) to insert the Luciferase gene into the pDonor221 donor vector according to the manufacturer’s protocol and create the Luciferase middle entry vector. The construct was transformed into competent Top10 E. coli cells, purified, and validated using a digest with Pvu II NEB). An LR reaction was carried out using Gateway LR Clonase II (Invitrogen) to insert the per2 5’ entry vector, the Luciferase middle entry vector, and the polyA 3’ entry vector (p3E-polyA, Tol2kit) into the destination vector pDestTol2pA according to the manufacturer’s instructions. The construct was transformed into Top10 E. coli cells, purified, and sent for full plasmid sequencing by Plasmidsaurus.

### Generation of 4xE-box:Luc and per2:Luc transgenic zebrafish lines

4xE-box:Luc^33^ constructs were a gift from Thomas Dickmeis and were validated by whole plasmid sequencing using Plasmidsaurus. The 4xE-box:Luc and per2:Luc transgenic lines were generated using the Tol2 system. Briefly, transposase mRNA was synthesized using the mMESSAGE mMACHINE SP6 Kit (Invitrogen). Immediately following the collection of fertilized eggs from wildtype (AB) adults, a mixture of 20 ng/µL transposase and 20 ng/µL 4xE-box:Luc or per2:Luc plasmid diluted in 0.5% phenol red solution (Fisher Scientific, Cat No. S93322) was injected into each egg at ∼2 nl per egg. Adult founders (F_0_) were out-crossed with wildtype (AB) adults to generate the F1 progeny. In-crossed F_1_ or F_2_ progeny were used for experiments.

### Generation of Crispants

Crispants knockouts were generated based on Kroll et alt. 2021^29^. Briefly, gRNAs targeting two regions of the genes *per2, cry1a, bmal1a, bmal1b*, and *bmal2*, were designed and ordered through IDT Atl-R^TM^ CRISPR-Cas9 system. crRNAs were annealed with tracrRNA to generate gRNAs, which were next mixed with Cas9 protein to generate a ribonucleoprotein (RNP) complex. One nanoliter of a mix of *per2/cry1a* RNPs or *bmal1a, bmal1b, bmal2* RNPs was injected into 1-cell stage zebrafish embryos using a microinjector. Dead and unfertilized fish were monitored and separated from fertilized eggs, which were screening for rhythms immediately after injection.

### Mutational Analysis

Zebrafish larvae were collected at the end of the screening period (7 dpf). The E3 media containing luciferin was discarded and fish were euthanized following UCSC Animal Protocol procedures. Single larvae were placed in 0.2 ml tubes and DNA was extracted using the HotShot Method^34^. For identification of indel mutations, each target region was amplified by PCR using Q5 High Fidelity Polymerase (NEB). The PCR product was then denatured and re-annealed, and heteroduplexes were digested with T7 Endonuclease I, that cleaves mismatched DNA, generating a fragmentation pattern in mutated samples.

### Developmental tracking of larvae - luminescence recordings

Adults were crossed overnight, and eggs were collected the next morning at lights on. The spawning of egg was determined as ZT0 and synchronized with lights on so that ZT0 corresponds with 0 hpf. Eggs were collected in 1X E3 media (5 mM NaCl, 0.17 mM KCl, 0.25 mM CaCl_2_, 0.16 mM MgSO_4_) with 1 mM D-Luciferin (eLUCK-1G / GoldBio) and placed with chorion in a white 96-well Optiplate^TM^ (Revvity) sealed with a Microseal B PCR plate film, to avoid media evaporation. Luminescence recordings were obtained every 30 minutes from 0 to 7 days using Tecan Spark plate reader on the kinetic settings with 12.5s integration time. Temperature was constant at 28°C. For 12h:12h light:dark exposure protocols, a 1500 lux white led lamp was positioned over the plate carrier and the plate was exposed to light outside the plate reader for 8 minutes, to then resume readings inside the machine. This was repeated every 30 minutes for 12 hours, to then continue readings in the dark for the next 12h. This loop was repeated for 7 days. For inverted cycles, constant light and light-induced phase shift protocols the same approach was used but the period of light exposure was changed accordingly. For constant dark reads, plates were maintained inside the machine.

### Rhythmicity and Period Tests

Oscillatory expression was analyzed in Biodare2 using the eJTK rhythmicity test for 24h period data or BD2-Classic for non-24h data, with significant values considered *p*<0.05. Period analysis was made using Fast Fourier Transform Non-Linear Least Squares (FFT-NLLS).

## Figure Legends

**Supplemental Figure 1. Free-run oscillations of *per3 and E-box* reporter are shifted**. Luminescence reads from *per3:luc* (blue line) and *4XE-box:luc* (grey line) zebrafish embryos entrained in 12h:12h LD cycles from 0 to 96 hpf and then maintained in free run (constant darkness) until 168 hpf. *Per3:luc* oscillates in LD and continues to cycle in constant darkness with a peak from the last light exposure (∼ZT6). *E-box:luc* shows no circadian oscillations in LD but gain rhythms after 96 hpf with a peak at ∼ZT24/0 RLU = Relative Luminescence Units. N= Number of animals per group.

**Supplemental Figure 2. The ribonucleoprotein injection process does not perturb the circadian clock**. Luminescence reads from *per3:luc* zebrafish embryos injected or not with a RNP complex of Cas9 protein and gRNAs targeting Tyrosinase, as injection control. Embryos were maintained in 12h:12h LD cycles from 0 to 168 hpf. No changes in *per3* rhythms are observed due to the injection process. RLU = Relative Luminescence Units. N= Number of animals per group.

## Acknowledgements

We would like to thank Thomas Dickmeis for generously sharing the 4xE-box:luc construct. We would like to thank undergraduate researchers Jared Kolahi, Emily Passion, and Eric Delgado, as well as Santiago Ruiz Solis for their assistance with maintaining the lines used here and general zebrafish facility care.

## References

1. Patke A, Young MW, Axelrod S. Molecular mechanisms and physiological importance of circadian rhythms. Nat Rev Mol Cell Biol. 2020;21(2):67–84. doi:10.1038/s41580-019-0179-2

2. Partch CL, Green CB, Takahashi JS. Molecular architecture of the mammalian circadian clock. Trends in Cell Biology. 2014;24(2):90–99. doi:10.1016/j.tcb.2013.07.002

3. Brown LS, Doyle FJ. A dual-feedback loop model of the mammalian circadian clock for multi-input control of circadian phase. Ermentrout B, ed. PLoS Comput Biol. 2020;16(11):e1008459. doi:10.1371/journal.pcbi.1008459

4. Landgraf D, Achten C, Dallmann F, Oster H. Embryonic development and maternal regulation of murine circadian clock function. Chronobiology International. 2015;32(3):416–427. doi:10.3109/07420528.2014.986576

5. Greiner P, Houdek P, Sládek M, Sumová A. Early rhythmicity in the fetal suprachiasmatic nuclei in response to maternal signals detected by omics approach. Kramer A, ed. PLoS Biol. 2022;20(5):e3001637. doi:10.1371/journal.pbio.3001637

6. Zhao J, Warman GR, Stanewsky R, Cheeseman JF. Development of the Molecular Circadian Clock and Its Light Sensitivity in Drosophila Melanogaster. J Biol Rhythms. 2019;34(3):272–282. doi:10.1177/0748730419836818

7. Delaunay F, Thisse C, Marchand O, Laudet V, Thisse B. An Inherited Functional Circadian Clock in Zebrafish Embryos. Science. 2000;289(5477):297–300. doi:10.1126/science.289.5477.297

8. Delaunay F, Thisse C, Thisse B, Laudet V. Differential regulation of Period 2 and Period 3 expression during development of the zebrafish circadian clock. Gene Expression Patterns. 2003;3(3):319–324. doi:10.1016/S1567-133(03)00050-4

9. Dekens MP, Whitmore D. Autonomous onset of the circadian clock in the zebrafish embryo. EMBO J. 2008;27(20):2757–2765. doi:10.1038/emboj.2008.183

10. Kaneko M, Cahill GM. Light-Dependent Development of Circadian Gene Expression in Transgenic Zebrafish. Schibler U, ed. PLoS Biol. 2005;3(2):e34. doi:10.1371/journal.pbio.0030034

11. Vatine G, Vallone D, Appelbaum L, et al. Light directs zebrafish period2 expression via conserved D and E boxes. PLoS Biol. 2009;7(10):e1000223. doi:10.1371/journal.pbio.1000223

12. Li Y, Li G, Wang H, Du J, Yan J. Analysis of a Gene Regulatory Cascade Mediating Circadian Rhythm in Zebrafish. Jensen LJ, ed. PLoS Comput Biol. 2013;9(2):e1002940. doi:10.1371/journal.pcbi.1002940

13. Ben-Moshe Z, Alon S, Mracek P, et al. The light-induced transcriptome of the zebrafish pineal gland reveals complex regulation of the circadian clockwork by light. Nucleic Acids Research. 2014;42(6):3750–3767. doi:10.1093/nar/gkt1359

14. Merrow M, Spoelstra K, Roenneberg T. The circadian cycle: daily rhythms from behaviour to genes: First in the Cycles Review Series. EMBO Reports. 2005;6(10):930–935. doi:10.1038/sj.embor.7400541

15. Vallone D, Gondi SB, Whitmore D, Foulkes NS. E-box function in a period gene repressed by light. PNAS. 2004;101(12):4106–4111. doi:10.1073/pnas.0305436101

16. Nakahata Y, Yoshida M, Takano A, et al. A direct repeat of E-box-like elements is required for cell-autonomous circadian rhythm of clock genes. BMC Molecular Biol. 2008;9(1):1. doi:10.1186/1471-2199-9-1

17. Mracek P, Santoriello C, Idda ML, et al. Regulation of per and cry Genes Reveals a Central Role for the D-Box Enhancer in Light-Dependent Gene Expression. PLOS ONE. 2012;7(12):e51278. doi:10.1371/journal.pone.0051278

18. Pagano C, Siauciunaite R, Idda ML, et al. Evolution shapes the responsiveness of the D-box enhancer element to light and reactive oxygen species in vertebrates. Sci Rep. 2018;8(1):13180. doi:10.1038/s41598-018-31570-8

19. Kwak JS, León-Tapia MÁ, Diblasi C, et al. Functional and regulatory diversification of Period genes responsible for circadian rhythm in vertebrates. McIntyre L, ed. G3: Genes, Genomes, Genetics. 2024;14(10):jkae162. doi:10.1093/g3journal/jkae162

20. Hirayama J, Alifu Y, Hamabe R, et al. The clock components Period2, Cryptochrome1a, and Cryptochrome2a function in establishing light-dependent behavioral rhythms and/or total activity levels in zebrafish. Sci Rep. 2019;9(1):196. doi:10.1038/s41598-018-37879-8

21. Wang M, Zhong Z, Zhong Y, Zhang W, Wang H. The Zebrafish Period2 Protein Positively Regulates the Circadian Clock through Mediation of Retinoic Acid Receptor (RAR)-related Orphan Receptor α (Rorα). Journal of Biological Chemistry. 2015;290(7):4367–4382. doi:10.1074/jbc.M114.605022

22. Ruggiero G, Ben-Moshe Livne Z, Wexler Y, et al. Period 2: A Regulator of Multiple Tissue-Specific Circadian Functions. Front Mol Neurosci. 2021;14:718387. doi:10.3389/fnmol.2021.718387

23. Yang M, Liu Y, Zhong Z, et al. Direct regulation of Per2 by Roraa: insights into circadian and metabolic interplay in zebrafish. Cell Mol Life Sci. 2025;82(1):195. doi:10.1007/s00018-025-05696-8

24. Katsuyama Y, Oomiya Y, Dekimoto H, et al. Expression of zebrafish ROR alpha gene in cerebellar-like structures. Developmental Dynamics. 2007;236(9):2694–2701. doi:10.1002/dvdy.21275

25. Flores MV, Hall C, Jury A, Crosier K, Crosier P. The zebrafish retinoid-related orphan receptor (ror) gene family. Gene Expression Patterns. 2007;7(5):535–543. doi:10.1016/j.modgep.2007.02.001

26. Hurd MW, Cahill GM. Entraining Signals Initiate Behavioral Circadian Rhythmicity in Larval Zebrafish. J Biol Rhythms. 2002;17(4):307–314. doi:10.1177/074873002129002618

27. Wang H, Yang Z, Li X, et al. Correction: Single-cell in vivo imaging of cellular circadian oscillators in zebrafish. PLoS Biol. 2021;19(8):e3001382. doi:10.1371/journal.pbio.3001382

28. Hoshijima K, Jurynec MJ, Klatt Shaw D, Jacobi AM, Behlke MA, Grunwald DJ. Highly Efficient CRISPR-Cas9-Based Methods for Generating Deletion Mutations and F0 Embryos that Lack Gene Function in Zebrafish. Developmental Cell. 2019;51(5):645-657.e4. doi:10.1016/j.devcel.2019.10.004

29. Kroll F, Powell GT, Ghosh M, et al. A simple and effective F0 knockout method for rapid screening of behaviour and other complex phenotypes. eLife. 2021;10:e59683. doi:10.7554/eLife.59683

30. Quick RE, Buck LD, Parab S, Tolbert ZR, Matsuoka RL. Highly Efficient Synthetic CRISPR RNA/Cas9-Based Mutagenesis for Rapid Cardiovascular Phenotypic Screening in F0 Zebrafish. Front Cell Dev Biol. 2021;9:735598. doi:10.3389/fcell.2021.735598

31. Ren F, Lin Q, Gong G, et al. Igf2bp3 maintains maternal RNA stability and ensures early embryo development in zebrafish. Commun Biol. 2020;3(1):94. doi:10.1038/s42003-020-0827-2

32. Moravec CE, Voit GC, Otterlee J, Pelegri F. Identification of maternal-effect genes in zebrafish using maternal crispants. Development. 2021;148(19):dev199536. doi:10.1242/dev.199536

33. Weger M, Weger BD, Diotel N, et al. Real-time in vivo monitoring of circadian E-box enhancer activity: a robust and sensitive zebrafish reporter line for developmental, chemical and neural biology of the circadian clock. Dev Biol. 2013;380(2):259–273. doi:10.1016/j.ydbio.2013.04.035

34. Meeker ND, Hutchinson SA, Ho L, Trede NS. Method for Isolation of PCR-Ready Genomic DNA from Zebrafish Tissues. BioTechniques. 2007;43(5):610–614. doi:10.2144/000112619

