## Supplemental Figures for "Longitudinal Tracking Reveals Developmental Transitions in Zebrafish Clock Gene Expression"

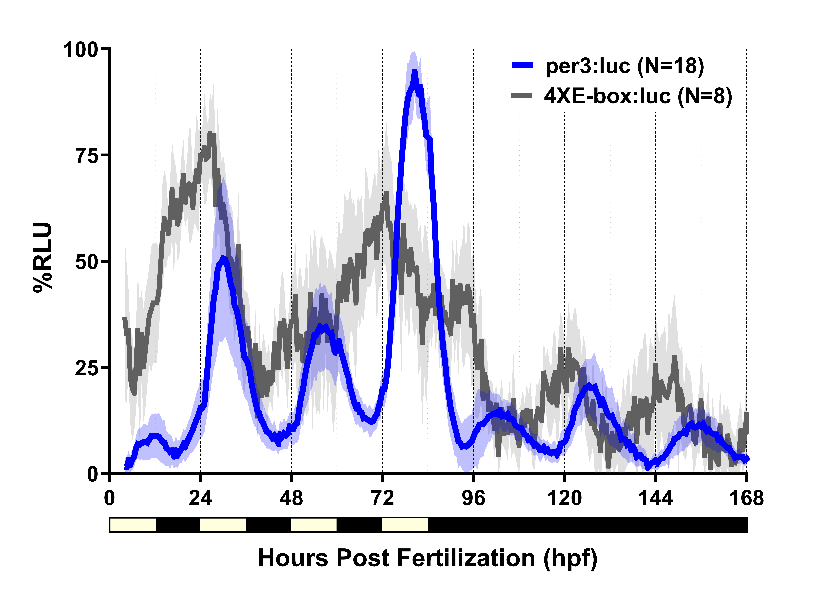

**Supplemental Figure 1. Free-run oscillations of *per3 and E-box* reporter are shifted.** Luminescence reads from *per3:luc* (blue line) and *4XE-box:luc* (grey line) zebrafish embryos entrained in 12h:12h LD cycles from 0 to 96 hpf and then maintained in free run (constant darkness) until 168 hpf**.**  *Per3:luc* oscillates in LD and continues to cycle in constant darkness with a peak from the last light exposure (~ZT6). *E-box:luc* shows no circadian oscillations in LD but gain rhythms after 96 hpf with a peak at ~ZT24/0 RLU = Relative Luminescence Units. N= Number of animals per group.

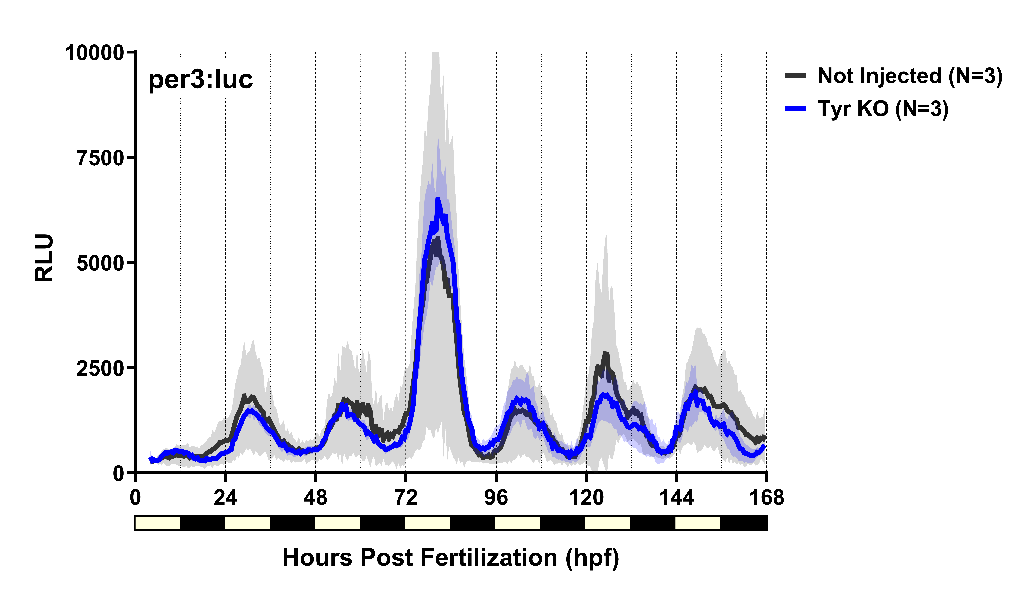

**Supplemental Figure 2. The ribonucleoprotein injection process does not perturb the circadian clock.** Luminescence reads from *per3:luc* zebrafish embryos injected or not with a RNP complex of Cas9 protein and gRNAs targeting Tyrosinase, as injection control. Embryos were maintained in 12h:12h LD cycles from 0 to 168 hpf. No changes in *per3* rhythms are observed due to the injection process. RLU = Relative Luminescence Units. N= Number of animals per group.

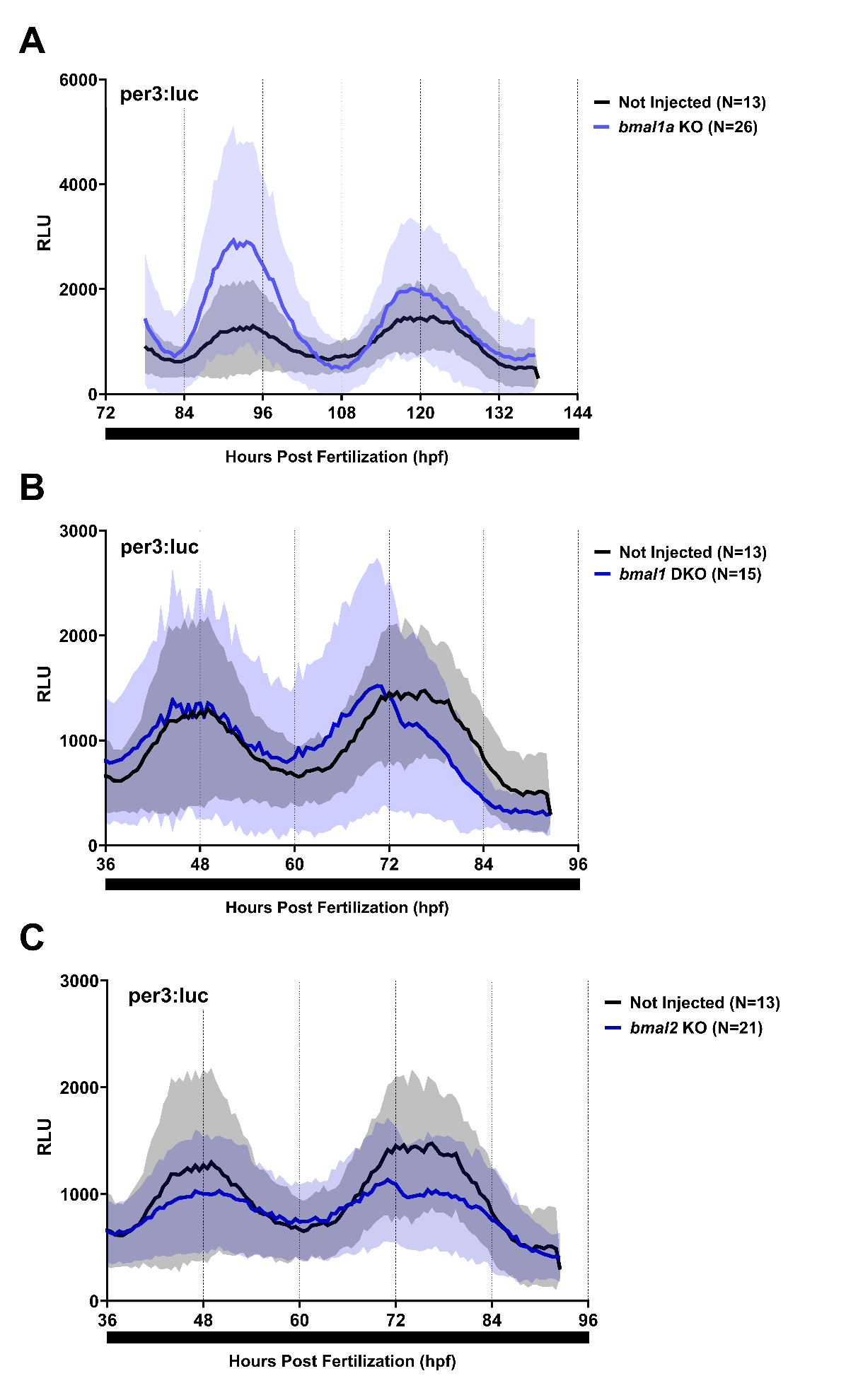

**Supplemental Figure 3. Zebrafish *bmal* paralogs show redundancy in maintaining *per3* rhythms.** Luminescence reads from *per3:luc* zebrafish embryos injected or not with a RNP complex of Cas9 protein and gRNAs targeting **(A)** *bmal1a*, **(B)** *bmal1a and bmal1b* (*bmal1* DKO) and **(C)** *bmal2* . Embryos were maintained in and incubator with 12h:12h LD cycles from 0 to 36 hpf and then placed to record rhythms in constant darkness. No significant changes in *per3* rhythms are observed in single or double *bmal* mutants. RLU = Relative Luminescence Units. N= Number of animals per group.
